# *PPD-H1* Improves Stress Resistance and Energy Metabolism to Boost Spike Fertility under High Ambient Temperatures

**DOI:** 10.1101/2024.11.04.621966

**Authors:** Tianyu Lan, Agatha Walla, Kumsal Ecem Çolpan Karışan, Gabriele Buchmann, Vera Wewer, Sabine Metzger, Isaia Vardanega, Einar Baldvin Haraldsson, Gesa Helmsorig, Venkatasubbu Thirulogachandar, Rüdiger Simon, Maria von Korff

## Abstract

High ambient temperature (HT) impairs reproductive development and grain yield in temperate crops. To ensure reproductive success under HT, plants must maintain developmental stability. However, the mechanisms integrating plant development and temperature resilience are largely unknown. Here, we demonstrate that *PHOTOPERIOD 1* (*PPD-H1*), homologous to *PSEUDO RESPONSE REGULATOR* genes of the Arabidopsis circadian clock, controls developmental stability in response to HT in barley. We analyzed HT responses in independent introgression lines with either the ancestral wild-type *Ppd-H1* allele or the natural *ppd-h1* variant, selected in spring varieties to delay flowering and enhance yield under favourable conditions. HT delayed inflorescence development and reduced grain number in *ppd-h1* mutant lines, while the wild-type *Ppd-H1* genotypes accelerated reproductive development and showed a stable grain set under HT. Using a CRISPR/Cas9-induced *ppd-h1* mutant, we confirmed that the CCT domain of *Ppd-H1* controls developmental stability, but not clock gene expression. Transcriptome and phytohormone analyses in developing leaves and inflorescences revealed increased stress gene expression and abscisic acid levels in the leaf and inflorescence of the natural and induced mutant *ppd-h1* lines. Furthermore, the mutant *ppd-h1* lines downregulated photosynthesis-and energy metabolism-related genes, and reduced auxin and cytokinin levels in the inflorescence, which impaired anther and pollen development. By contrast, in the wild-type *Ppd-H1* plants, the transcriptome and phytohormone levels and anther and pollen development remained stable under HT. Our findings suggest that *Ppd-H1* enhances stress resistance and energy metabolism, thereby stabilizing reproductive development, floret fertility and grain set under HT.

## Introduction

An increase in the global average temperature has strong adverse effects on crop yield (Tashiro and Wardlaw, 1989; Hakala *et al*., 2011; Ottman *et al*., 2012; Asseng *et al*., 2015; Xie *et al*., 2018). In particular, reproductive development, which is critical for determining the number of spikelets, fertile florets, and grains, is very susceptible to high temperatures in the temperate crops barley and wheat (Jacott and Boden, 2020). Understanding the genetic, molecular, and metabolic bases for high temperature-mediated changes in reproductive development are crucial for breeding thermotolerant barley and wheat varieties.

High ambient temperature (HT) below the heat-stress threshold induces a suite of morphological and architectural changes in plants. These processes are collectively referred to as thermomorphogenesis (Delker, Quint and Wigge, 2022). In the model plant *Arabidopsis thaliana*, HT accelerates flowering and overcomes the short-day (SD)-induced delay in flowering through upregulation of the florigen *FLOWERING LOCUS T* (*FT*) in Arabidopsis (Balasubramanian 2006). HT causes hyponasty and enhances the elongation of hypocotyl, stem, and leaf petiole, but reduces leaf blade size, which improves the transpirational cooling capacity of leaves and shoots (Casal and Balasubramanian, 2019). The *BASIC HELIX–LOOP–HELIX* (*bHLH*) transcription factor *PHYTOCHROME INTERACTING FACTOR 4* (*PIF4*) plays a key role in thermomorphogenesis by orchestrating transcriptional changes that trigger *FT* transcription and flowering, as well as the phytohormone-induced hypocotyl and petiole elongation responses (Koini *et al*., 2009; Franklin *et al*., 2011; Kumar *et al*., 2012). Variation in ambient temperature is sensed by the photoreceptor PHYTOCHROME B (PhyB). Warm temperatures accelerate the thermal/dark reversion of PhyB from its active to inactive form, thereby relieving the repression of the *PIF4* (Jung *et al*., 2016; Legris *et al*., 2016; Delker, van Zanten and Quint, 2017). In addition, another putative thermosensor, the circadian clock gene *EARLY FLOWERING 3* (*ELF3*) integrates temperature signals to the circadian oscillator, thereby coordinating the rhythmic control of thermoresponsive physiological outputs (Ezer *et al*., 2017; Jung *et al*., 2020; Zhu, Quint and Anwer, 2022). Furthermore, epigenetic modifications and chromatin remodelling play a prominent role in thermomorphogenesis, where HT decreases histone occupancy of DNA, thereby improving DNA accessibility and transcriptional activation or repression of thermomorphogenic genes and *FT* (Kumar *et al*., 2012; Huai *et al*., 2018; Van Der Woude *et al*., 2019).

While many of the phenotypic and molecular responses to HT are conserved across species, the temperate cereal crops barley and wheat display ambient temperature responses distinct from those observed in Arabidopsis. In barley and wheat, HT reduces stem elongation, and plants become more compact with reduced plant height and total biomass (Batts *et al*., 1997; Hemming *et al*., 2012; Dixon *et al*., 2019). Furthermore, HT displays genotype-specific effects on reproductive development, either accelerating or delaying flowering time in both species. Recent studies have identified the photoperiod response gene, *PHOTOPERIOD 1* (*PPD-H1*), a barley homolog of Arabidopsis *PSEUDO RESPONSE REGULATOR* (*PRR*) genes from the circadian clock, as a major regulator of thermo-responsive flowering in barley (Ejaz and von Korff, 2017; Ochagavía *et al*., 2022). Originally, the ancestral dominant *Ppd-H1*, hereafter referred to as the wild-type *Ppd-H1* allele, was identified as a photoperiod response gene that accelerates flowering under inductive long-day conditions (LD) by upregulating *FLOWERING LOCUS T1* (*FT1*), a homolog of Arabidopsis *FT*, in the leaf (Turner *et al*., 2005; Digel, Pankin and von Korff, 2015). A natural mutation in the CONSTANS, CO-like, and TOC1 (CCT) domain of *Ppd-H1*, hereafter referred to as mutant *ppd-h1* allele, is prevalent in spring barley causes a reduction in *FT1* expression and a delay in flowering time under LD (Turner *et al*., 2005; Digel, Pankin and von Korff, 2015). Moreover, *PPD-H1* also controls flowering time in response to HT, where the spring barley cultivars with mutant *ppd-h1* allele show a delay in flowering time under HT, but the derived near-isogenic lines (NILs) with wild-type *Ppd-H1* alleles display accelerated flowering under HT (Ejaz and von Korff, 2017). This allele-specific delay in flowering time under HT was linked to reduced floret fertility and grain number per spike. By contrast, genotypes with the wild-type *Ppd-H1* allele maintained high floret fertility and grain set under HT (Ejaz and von Korff, 2017). In contrast to Arabidopsis, where *FT* was strongly upregulated by HT (Balasubramanian 2006), transcript levels of *FT1* were always downregulated by HT in the leaf and also in the NILs with the wild-type *Ppd-H1* allele that accelerated flowering under HT (Ejaz and von Korff, 2017). The *PPD-H1*-dependent developmental timing in response to temperature is thus likely mediated by developmental regulators other than *FT1* in barley.

The aim of this study was to dissect the effects of HT and *PPD-H1* on plant growth, inflorescence meristem (IM) activity, spikelet induction rate, floret development, and spike fertility. Furthermore, our objective was to detect genes and regulatory networks in the leaf, shoot apex, and inflorescence that underly the *PPD-H1*-dependent developmental responses to HT. We demonstrate that *PPD-H1* interacts with ambient temperature to control IM activity, thereby influencing the rate and duration of spikelet meristem (SM) and floral meristem (FM) induction, and floret fertility. Through global transcriptome and phytohormone analyses, we demonstrate that *PPD-H1* plays a crucial role in stress resistance under HT. Differential expression of genes involved in stress response in the leaf and photosynthesis and energy metabolism in the inflorescence suggests that wild-type *Ppd-H1* enhances stress resistance, thereby maintaining energy supply to drive reproductive development and mitigate the adverse effects of HT on floret fertility.

## Results

### High Ambient Temperature Affects Plant Development

To evaluate the interaction between ambient temperatures and *PPD-H1* on plant growth and development, we characterized shoot and inflorescence development in two independent pairs of near-isogenic lines (NILs), differing for a natural mutation in the CCT domain of *PPD-H1* and one CRISPR/Cas-induced *ppd-h1* mutant line under control (CT, 20 °C/16 °C, day/night) and high ambient temperature (HT, 28 °C/24 °C, day/night) in inductive long-day (LD, 16 h/8 h, light/dark) conditions. We focused our analyses on the spring barley cultivar Golden Promise (GP), which carries a natural mutation in the CCT domain of the ancestral wild-type *Ppd-H1*, causing a Gly-to-Trp substitution at 657 AA (Supplementary Dataset 1) and the derived NIL Golden Promise-fast (GP-fast) with a wild-type *Ppd-H1* allele from the winter barley cultivar Igri. We further generated a CRISPR/Cas9-induced *Ppd-H1* mutant (*ppd-h1.1*) in the GP-fast background, which carried a 1-bp insertion upstream of the CCT domain, resulting in a truncated *PPD-H1* protein without the CCT domain (Supplementary Fig. S1, Supplementary Dataset 1). The induced mutant *ppd-h1.1* did not show differences in diurnal expression of major clock genes as already demonstrated for the natural *ppd-h1* mutants Scarlett and GP (Campoli et al., 2012) (Supplementary Fig. S2). To verify the effects of *PPD-H1* on developmental, morphological, and molecular responses, we used an independent pair of NILs, the spring barley cultivar Scarlett (*ppd-h1*) and the derived NIL S42-IL107 (*Ppd-H1*) that carry the same putatively functional amino acid variant in the CCT domain as GP and GP-fast (Supplementary Dataset 1) and have been scored for developmental responses to HT by Ejaz et al. (2017).

Under CT, genotypes with a mutant *ppd-h1* allele, GP and *ppd-h1.1*, exhibited a delay in flowering time compared to GP-fast with a wild-type *Ppd-H1* allele (Fig. 1A, B). HT further delayed flowering time in GP and *ppd-h1.1* by an average of ten days but accelerated flowering time in GP-fast by an average of four days compared to CT. HT thus interacted with *PPD-H1* to control flowering time (Fig. 1B), as reported for Scarlett and S42-IL107 (Ejaz and von Korff, 2017). We further investigated the impact of HT on inflorescence development by dissecting the main shoot apex (MSA) of GP-fast, GP, and *ppd-h1.1* every three to five days. MSA development from seedling emergence to pollination was evaluated according to the Waddington scale (Waddington et al. 1983). The Waddington scale assesses developmental stages by examining the morphogenesis of the shoot apex and the carpel of the most advanced floret of the main spike. During vegetative development, leaf primordia are initiated until the induction of spikelet meristems (SMs), which become visible at the double ridge stage (W2.0). The differentiation of the first floral meristem (FM) and stem elongation start at the stamen primordium stage (W3.5). At W6.0, all floret organ primordia are initiated, followed by rapid exponential spike growth and floral organ maturation from W7.0 to W10.0, when anthesis and pollination take place. In all genotypes, HT caused a significant delay in the transition from the vegetative to spikelet initiation stage (W2.0) (Fig. 1C). However, the spikelet induction phase (W2.0-W3.5) and floral development (W3.5-W10.0) were delayed by HT in GP and *ppd-h1.1* but accelerated in GP-fast compared to CT (Fig. 1C). The *PPD-H1*-dependent effects of HT on floral development were thus responsible for the observed delay in flowering time in GP and *ppd-h1.1* and the earlier flowering time in GP-fast. The number of tillers and spikes were increased under HT in the late flowering *ppd-h1* mutant plants, GP, Scarlett, and *ppd-h1.1*, but were not affected in the wild-type *Ppd-H1* plants GP-fast and S42-IL107 (Fig.1D, E, Supplementary Fig. S3A, B, D). By contrast, plant height was strongly reduced by HT in all genotypes (Fig. 1F, Supplementary Fig. S3C). The leaf appearance rate (LAR) was accelerated under HT in both GP and GP-fast compared to CT (Supplementary Fig. S4A). Additionally, HT altered leaf size and form; most leaves became longer and narrower, while the last leaf, the flag leaf, was shorter and narrower, and these effects were more pronounced in GP than GP-fast (Supplementary Fig. S4B). Analysis of the anatomy of the second leaf (L2) suggested that the changes in leaf shape might be due to a strong reduction in cell size and an increase in cell number under HT compared to CT (Supplementary Fig. S4C-E). Leaf senescence was strongly enhanced by HT in GP and *ppd-h1.1* but not altered in GP-fast based on the percentage of the non-green area of L2 on the main culm (Supplementary Fig. S4F). Together, HT interacted with *PPD-H1* to control inflorescence development, flowering time, tiller and spike numbers, and leaf senescence rate. By contrast, HT reduced plant height, accelerated LAR, and altered leaf form independent of *PPD-H1*.

**Figure 1.**
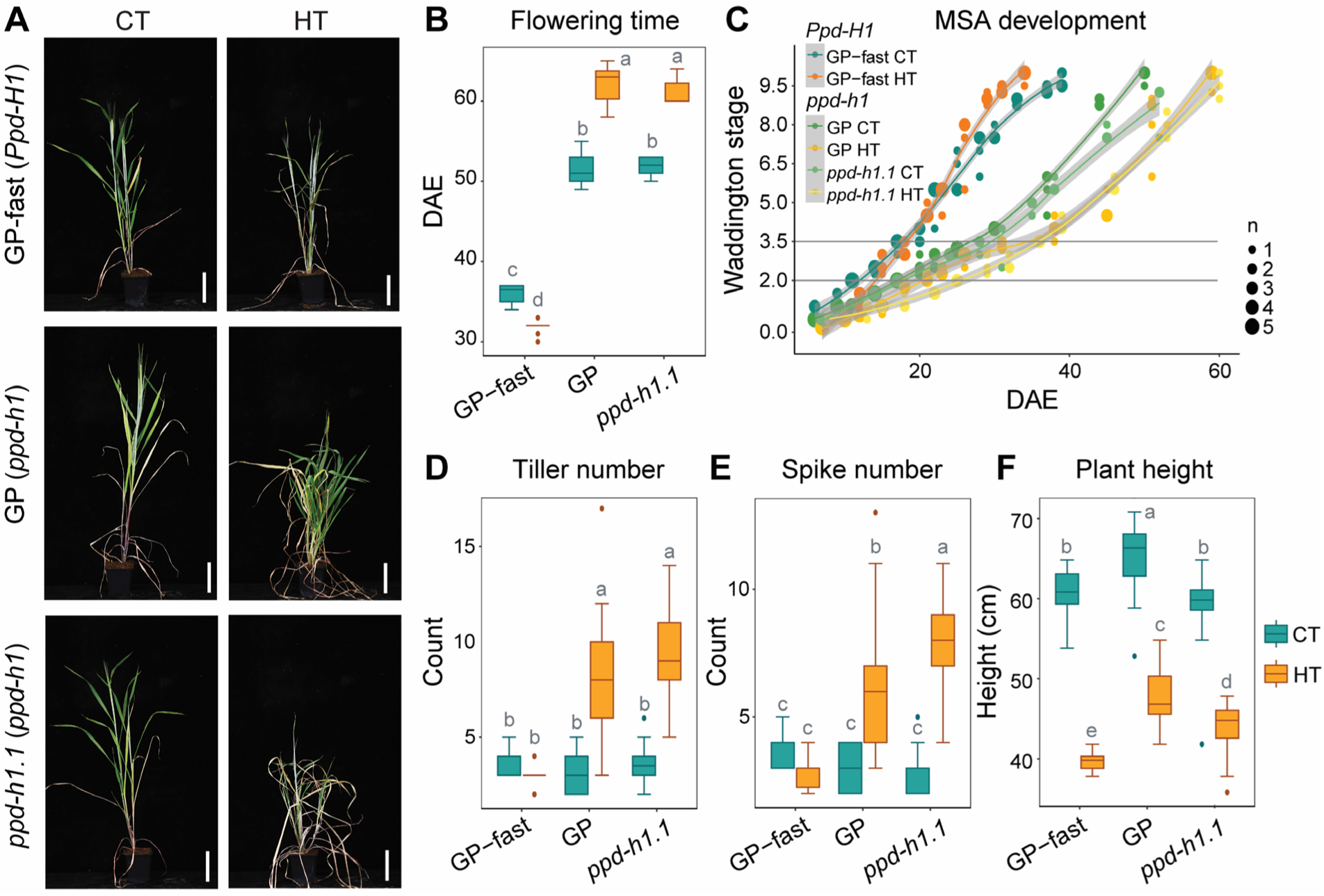
Effects of High Ambient Temperature and *PPD-H1* on Plant Growth and Spike Development. (A) Representative plants of spring barley cultivar Golden Promise (GP, *ppd-h1*), its derived near-isogenic line GP-fast (*Ppd-H1*), and CRISPR/Cas9-induced *ppd-h1* mutant *ppd-h1.1* (*ppd-h1*) under control (CT, 20 °C/16 °C, day/nght) and high ambient temperatures (HT, 28 °C/24 °C, day/night) at 55 days after emergence (DAE). Scale bar = 10 cm. (B) Comparison of flowering time (in days from plant emergence to awn appearance) among GP-fast (*Ppd-H1*), GP (*ppd-h1*), and *ppd-h1.1* (*ppd-h1*) under CT and HT. n ≥ 15. (C) Development of the main shoot apex (MSA) of GP-fast (*Ppd-H1*), GP (*ppd-h1*), and *ppd-h1.1* (*ppd-h1*) from emergence to postpollination under CT and HT, according to Waddington et al. (1983). The horizontal lines indicate the spikelet induction stage (W2.0) and stamen primordium stage (W3.5). n = 3-5. Dot sizes indicate the number of overlapping data points. The trend line is calculated using a polynomial regression (loess smooth line), and the grey area shows the 95% confidence interval. (D-F) Comparison of the number of tillers (D) and spikes (E), and plant height (F) among GP-fast (*Ppd-H1*), GP (*ppd-h1*), and *ppd-h1.1* (*ppd-h1*) at maturity under CT and HT. Each box shows the median and interquartile range (IQR), and whisker lines extend to the smallest and largest values within 1.5*IQR from the lower and upper quartiles, respectively. Outliers beyond this range are represented by individual points. Statistical groups were assigned using ANOVA followed by a Tukey’s post-hoc test. Different letters above the boxplots indicate significant differences between groups (*p* < 0.05). n ≥ 15.

### Ambient Temperature Interacts with *PPD-H1* to Regulate Spikelet Meristem Induction and Spike Fertility

Upon transition to reproductive growth, the indeterminate barley inflorescence meristem (IM) generates SMs on its flanks. In two-rowed barley GP and Scarlett, the central SM develops a single FM, which later develops into a floret and a grain. The duration of the spikelet induction phase is generally positively correlated with the final spikelet number (Digel, Pankin and von Korff, 2015). Accordingly, under CT, the late flowering *ppd-h1* genotypes, GP, *ppd-h1.1*, and Scarlett, produced more SMs, FMs, and florets on the MSA compared to the early-flowering genotypes GP-fast and S42-IL107 (Fig. 2A-C, Supplementary Fig. S5, Supplementary Table S1). In GP-fast and S42-IL107, HT further accelerated reproductive development and thus shortened the SM and FM induction phases and reduced SM, FM, and floret numbers compared to CT (Fig. 2A-C, Supplementary Fig. S5, Supplementary Table S1). However, in the mutant *ppd-h1* genotypes, HT extended the reproductive phase further but decreased SM, FM, and floret numbers compared to CT (Fig. 2A-C, Supplementary Fig. S5, Supplementary Table S1). *PPD-H1*-and temperature-dependent changes in the SM and FM induction rates contributed to differences in final SM numbers. During the primary SM induction phase (W2.0-4.5), HT increased the rate of SM induction in the wild-type *Ppd-H1* genotypes, from 1.3 to 1.6 SMs/day in GP-fast and from 1.9 to 2.1 SMs/day in S42-IL107, respectively (Fig. 2A, Supplementary Fig. S5, Supplementary Table S1). In contrast, HT decreased the SM induction rates in mutant *ppd-h1* genotypes from 1.6 to 0.8 SMs/day in GP, from 1.7 to 1.3 SMs/day in *ppd-h1.1*, and from 1.4 to 1.1 SMs/day in Scarlett (Fig. 2A, Supplementary Fig. S5, Supplementary Table S1). The SM induction rates were positively associated with a reduction in IM width and height in GP and GP-fast, respectively (Supplementary Fig. S6). These findings indicated that HT interacted with *PPD-H1* to affect IM activity and maintenance and, thus, SM induction rate. As observed for the SM induction rate, HT increased the FM induction rate in the wild-type *Ppd-H1* genotypes but slowed it down in the mutant *ppd-h1* genotypes (Fig. 2A, Supplementary Fig. S5, Supplementary Table S1).

**Figure 2.**
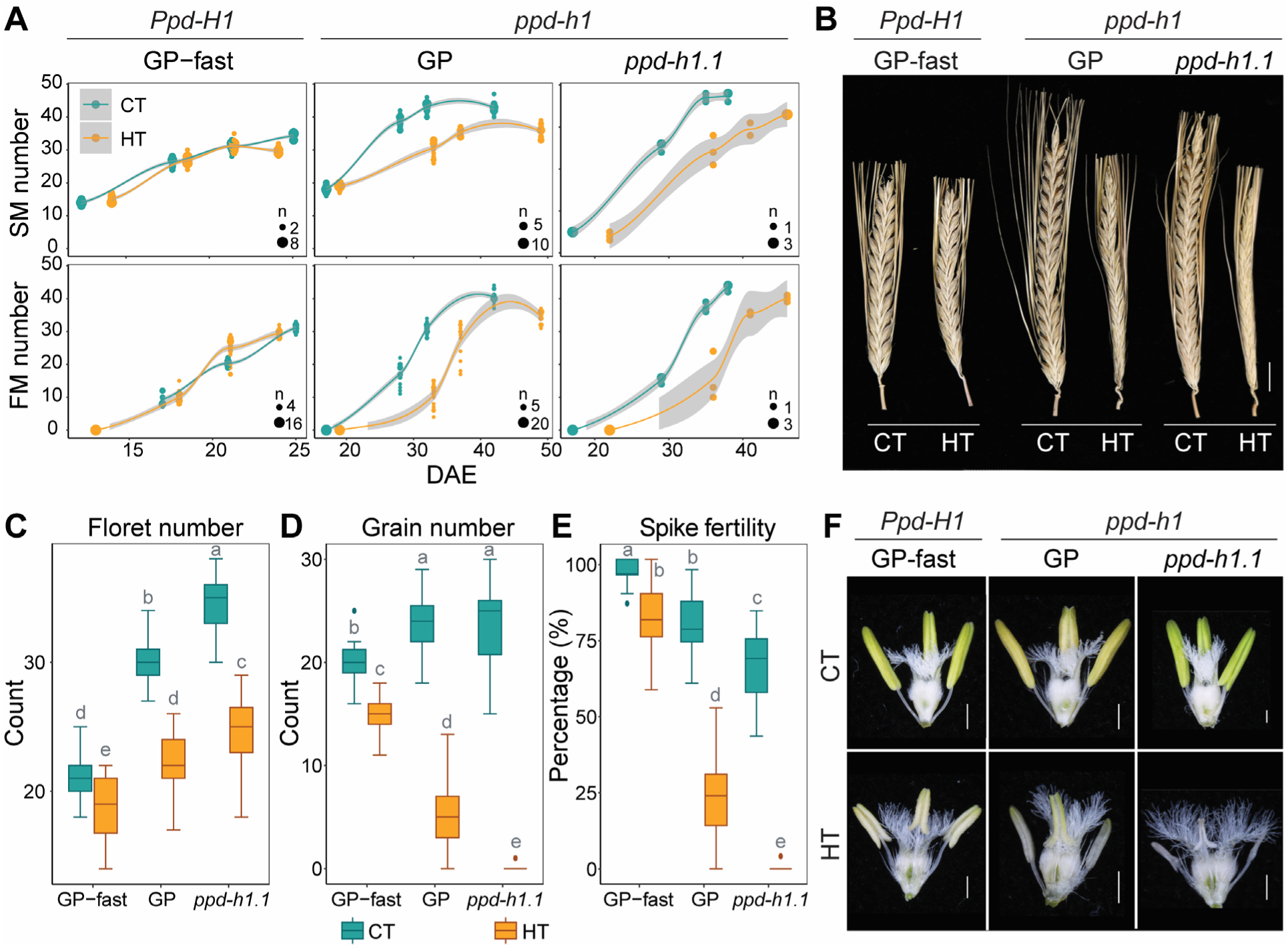
Effects of High Ambient Temperature on Reproductive Development. (A) Comparison of the number of spikelet meristems (SMs) and floral meristem (FMs) at spikelet induction (W2.0), stamen primordium (W3.5), carpel primordium (W4.5), and style primordium (W6.0) stages between control (CT, 20 °C/16 °C, day/night) and high ambient temperatures (HT, 28 °C/24 °C, day/night) in spring barley cultivar Golden Promise (GP, *ppd-h1*), its derived near-isogenic line GP-fast (*Ppd-H1*), and CRISPR/Cas9-induced mutant *ppd-h1.1* (*ppd-h1*). The x-axis indicates the corresponding days after emergence (DAE) to the Waddington stages W2.0-W6.0 (left to right). Dot sizes indicate the number of overlapping data points. Trend lines are calculated using a polynomial regression (loess smooth line), and the grey areas show a 95% confidence interval. n = 3-23. (B) Representative mature spikes on the main culm of GP-fast (*Ppd-H1*), GP (*ppd-h1*), and *ppd-h1.1* (*ppd-h1*) grown under CT and HT. Scale bar = 1 cm. (C-E) Comparison of the number of florets (C), grains (D), and spike fertility (E) at maturity on the main spike of GP-fast (*Ppd-H1*), GP (*ppd-h1*), and *ppd-h1.1* (*ppd-h1*) grown in CT and HT. Spike fertility is shown as the ratio of grain number to final floret number on the main culm. Each box shows the median and interquartile range (IQR), and whisker lines extend to the smallest and largest values within1.5*IQR from the lower and upper quartiles, respectively. Outliers beyond this range are represented by individual points. Statistical groups were assigned using ANOVA followed by Tukey’s post-hoc test. Different letters above the boxplots indicate significant differences between groups (*p* < 0.05). n ≥ 15. (F) Representative anthers and ovaries of GP-fast (*Ppd-H1*), GP (*ppd-h1*), and *ppd-h1.1* (*ppd-h1*) at W9.5 grown under CT and HT. Scale bar = 1 mm.

Although the final SM, FM, and floret numbers on the main spike were higher in the mutant *ppd-h1* compared to the wild-type *Ppd-H1* plants under both CT and HT, the final grain number was higher in the wild-type compared to mutant plants under HT (Fig. 2B-D, Supplementary Fig. S7A-C). Under CT, floret fertility, the number of grains produced per floret on the MSA, was 90-100% in the wild-type *Ppd-H1* and 70-80% in the mutant *ppd-h1* genotypes (Supplementary Fig. S7D, Supplementary Table S1). Under HT, floret fertility dropped to 80% in the wild-type *Ppd-H1* genotypes, but to 0-20% in mutant genotypes (Supplementary Table S1). HT caused strong floret abortion at all rachis nodes in the mutant *ppd-h1* genotypes, particularly at the base and tip of the spike, resulting in a strong reduction in grain number per spike under HT (Supplementary Fig. S7E, F). Impaired anther development in the *ppd-h1* mutant genotypes, as seen by the small, pale, and empty anthers in all florets along the spike, likely contributed to the reduction in floret fertility under HT (Fig. 2F). Although the anther size of GP-fast was also reduced under HT similar to GP and *ppd-h1.1*, pollen viability was only strongly reduced in the mutant but not wild-type plants (Fig. 2F, Supplementary Fig. S7G-J). In addition, HT altered the grain shape, leading to longer but narrower grains and a reduction in thousand-grain weight (TGW) independent of *PPD-H1* (Supplementary Fig. S7K-M).

In summary, *PPD-H1* interacted with ambient temperature to control IM activity and, thus, SM number and the rate of SM induction and floral development. Notably, the wild-type *Ppd-H1* genotypes enhanced floret fertility at basal and apical florets not only under HT but also under CT, possibly by regulating anther and pollen development.

### *PPD-H1* Controls Transcriptome and Phytohormone Variation in Response to High Ambient Temperature

To identify molecular changes responsible for the *PPD-H1*-dependent differences in development, floret fertility, and grain set in response to HT, we investigated global transcriptome changes in the leaf and MSA at different developmental stages under CT and HT. We collected the youngest fully elongated leaf and developing MSA of GP and GP-fast under CT and HT conditions at four developmental stages: vegetative (W1.0), spikelet induction (W2.0), stamen primordium (W3.5), and style primordium (W6.0) (Fig. 3A). To confirm the major effects of *PPD-H1* and HT on leaf and MSA transcriptomes, we harvested the same tissues for *ppd-h1.1* at the stamen primordium stage (W3.5), and Scarlett and S42-IL107 at the stamen primordium (W3.5) and style primordium (W6.0) stages under both temperature conditions.

**Figure 3.**
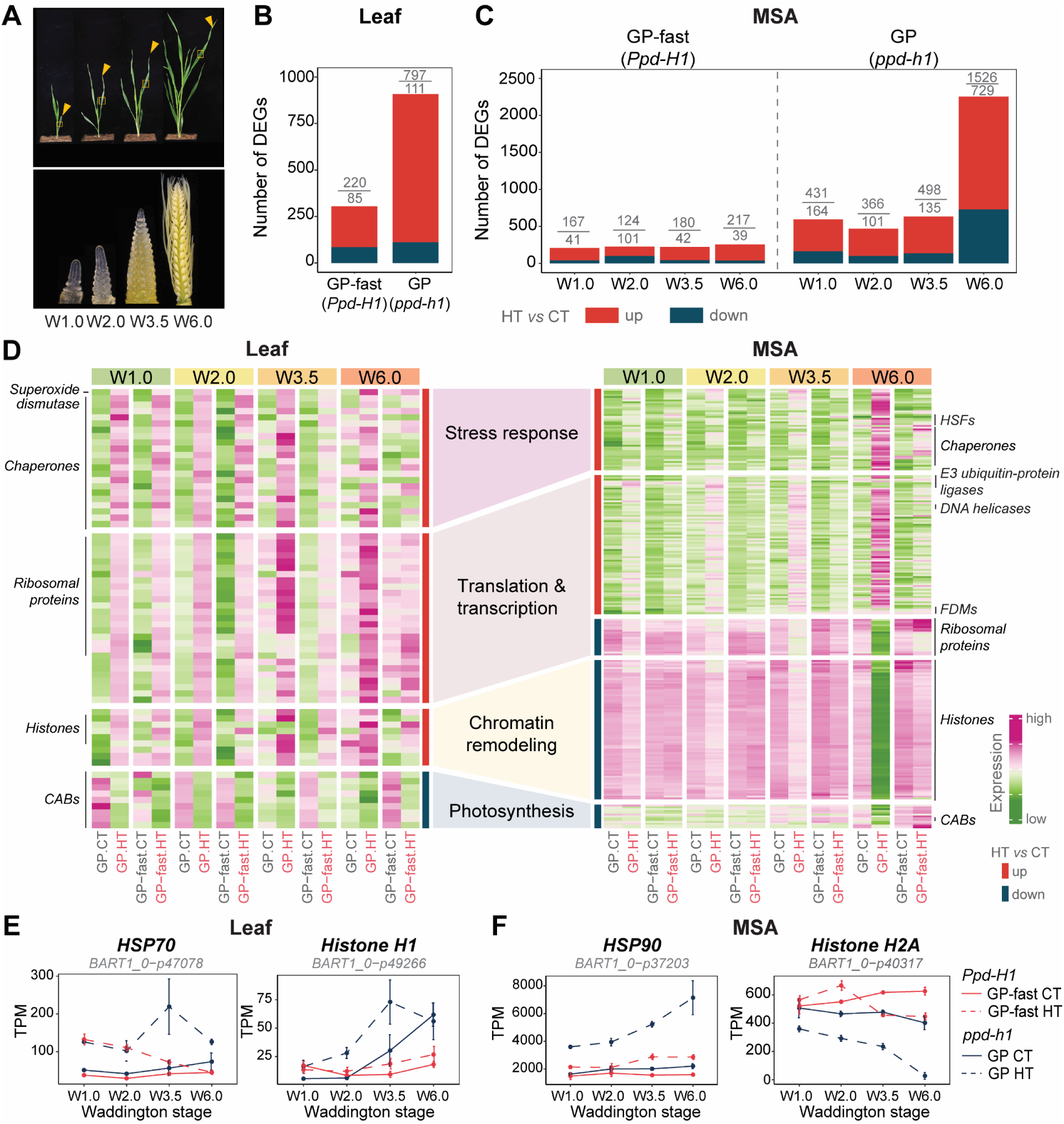
Thermal-Transcriptome Reprogramming in the Leaf and Shoot Apex. (A) Sampling strategy for studying transcriptomic changes in response to high ambient temperature (HT, 28 °C/24 °C, day/night) in the leaf and main shoot apex (MSA) at vegetative (W1.0), spikelet induction (W2.0), stamen primordium (W3.5), and style primordium (W6.0) stages. Orange arrows indicate the latest fully developed leaf on the main culm, from which the middle section was harvested for RNA-sequencing. Main shoot apices (MSAs) were collected at the same time as the leaf sample. (B) Number of differentially expressed genes (DEGs) that were upregulated (red) or downregulated (dark blue) in response to HT across at least three developmental stages in the leaf of Golden Promise (GP, *ppd-h1*) and its derived near-isogenic line GP-fast (*Ppd-H1*). |log_2_FC| ≥ 1, BH.FDR < 0.01. (C) Number of DEGs in MSA sample at each developmental stage in genotypes carrying different *PPD-H1* variants. |log_2_FC| ≥ 1, BH.FDR < 0.01. (D) Heatmaps represent the expression pattern of DEGs belonging to the top enriched gene ontology (GO) terms in the leaf (left) and MSA (right) of GP (*ppd-h1*) and GP-fast (*Ppd-H1*) at different developmental stages. The colour scale represents the range of Z-score normalized mean transcript per million (TPM) values of three to four biological replicates. The darker green hues represent lower expression, while darker magenta hues represent higher expression relative to other samples in the same gene. The representative gene families are labelled on the side of the heatmap. (E-F) Transcript profiles of representative genes, which are involved in heat response and chromatin remodelling in the leaf (E) and MSA (F) of GP-fast (*Ppd-H1*) (red) and GP (*ppd-h1*) (dark blue) grown under control ambient temperature (CT, 20 °C/16 °C, day/night) and HT. Transcript levels are shown in TPM. Solid and dashed lines indicate CT and HT, respectively. Error bars indicate the standard deviation of the mean of three to four biological replicates. *CAB*, *CHLOROPHYLL A-B BINDING*; *HSF*, *HEAT SHOCK FACTOR*; *FDM*, *FACTOR OF DNA METHYLATION*; *HSP70*, *HEAT SHOCK PROTEIN 70*; *HSP90, HEAT SHOCK PROTEIN 90.* For the gene filtering and statistics, see Supplementary Datasets 2 and 3.

The principal component analysis (PCA) revealed that temperature differences explained most of the variation in the leaf transcriptomes, followed by the genotype, while differences between stages were temperature and genotype-specific (Supplementary Fig. S8A, B). MSA samples showed a clear separation by the developmental stage, particularly for the samples at W6.0 (Supplementary Fig. S8C, D). The temperature only clearly differentiated the MSA transcriptomes of mutant *ppd-h1* genotypes but not the wild types (Supplementary Fig. S8C, D). To identify the significant effects of temperature, genotype, and developmental stage on transcript levels, we conducted a three-way ANOVA followed by pos-thoc pairwise comparisons between temperatures (HT versus CT) per stage and genotype in the leaf and MSA samples. Since leaf samples did not show a clear separation by developmental stages, we selected genes which were consistently regulated by temperature (|log_2_FC| ≥ 1, FDR < 0.01) across at least three developmental stages in GP-fast and GP and confirmed their temperature responsiveness in S42-IL107 and *ppd-h1.1* or Scarlett, respectively (Supplementary Fig. S9A, B). We thus detected 305 differentially expressed genes (DEGs) in GP-fast and 908 DEGs in GP confirmed in the other genotypes (Fig. 3B, Supplementary Fig. S9B, Supplementary Dataset 2). Since MSA samples were separated by stage, we identified DEGs for each stage separately (Supplementary Fig. S9C). In GP-fast, the number of DEGs increased from 208 DEGs at the vegetative stage (W1.0) to 256 DEGs at the floral development (W6.0) and thus remained relatively stable over the stages (Fig. 3C). The number of DEGs was considerably higher in GP at all stages with 595, 476, 633, and 2255 DEGs at W1.0, W2.0, W3.5, and W6.0, respectively (Figure 3C, Supplementary Fig. S9C-E). We focused our further analyses on those DEGs in the MSA, which were detected in at least two stages and/or confirmed in the independent *PPD-H1* variants (Fig. 3C, Supplementary Datasets 2, 3).

Gene ontology (GO) analysis revealed that in both leaf and MSA, DEGs upregulated under HT were enriched in stress response and transcriptional and translational regulation, while downregulated genes had functions in the light and dark reactions of photosynthesis (Fig. 3D, Supplementary Fig. S10, Supplementary Dataset 4). For instance, molecular chaperones like *HEAT SHOCK PROTEINs* (*HSPs*), *HEAT SHOCK TRANSCRIPTION FACTORs* (*HSFs*), and reactive oxygen species (ROS) scavengers were upregulated by HT in the leaf and MSA (Fig. 3D-F). By contrast, histone variants and chromatin/histone remodelers like *HISTONE* superfamily genes were upregulated in response to HT in the leaf but downregulated in the MSA (Fig.3D-F). Notably, transcriptional changes in the majority of these genes were more pronounced in the mutant *ppd-h1* than in the wild-type *Ppd-H1* genotypes; half of the temperature-responsive genes displayed significant genotype by temperature interaction effects (Fig. 3D, Supplementary Datasets 2, 3). The stronger transcriptional and phenotypic plasticity, leaf senescence, and floret abortion in mutant *ppd-h1* genotypes under HT suggested that wild-type *Ppd-H1* controls stress resistance. Indeed, under both CT and HT, genes with roles in biotic and abiotic stress responses were upregulated in the leaf, while the MSA genes involved in photosynthesis and energy metabolism were downregulated in the mutant compared to wild-type plants (GP-fast versus GP and *ppd-h1.1*, and S42-IL107 versus Scarlett) (Fig. 4A, B, Supplementary Dataset 4).

**Figure 4.**
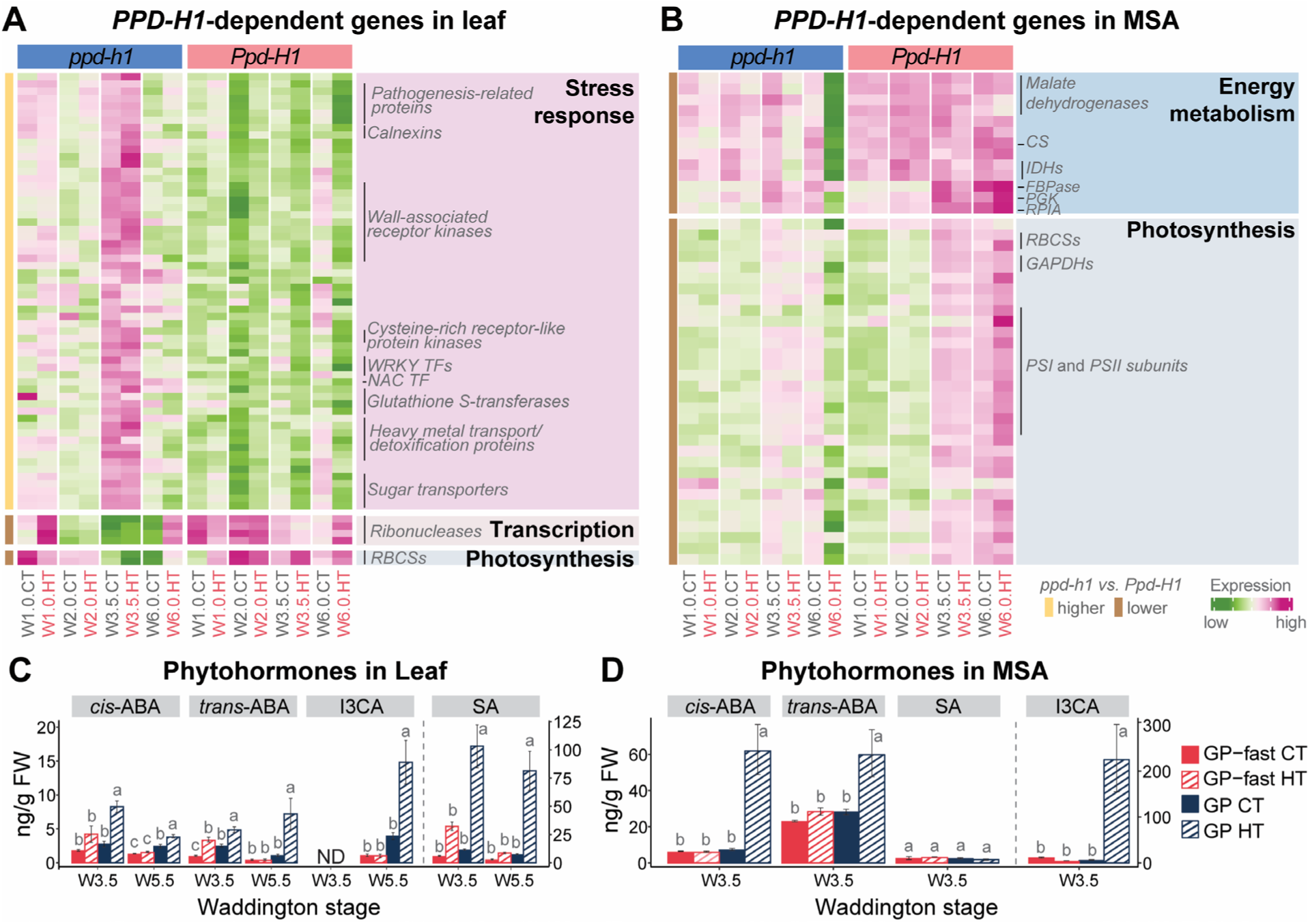
*PPD-H1*-Dependent Gene Identification and Phytohormone Changes in the Leaf and Shoot Apex. (A-B) Heatmaps display the expression pattern of *PPD-H1*-regulated genes from the enriched top gene ontology (GO) terms in the leaf (left) and MSA (right) of spring barley cultivar Golden Promise (GP, *ppd-h1*) and its near-isogenic line GP-fast (*Ppd-H1*) at different developmental stages and ambient temperatures. The colour scale represents the range of Z-score normalized mean transcript per million (TPM) values of three to four biological replicates. The darker green hues represent lower expression, while darker pink hues represent higher expression relative to other samples in the same gene. Representative gene families are labelled on the side of the heatmap. (C-D) Levels of the stress-related phytohormones, abscisic acid (ABA) and salicylic acid (SA), and the defense-related indolic compound, indole-3-carboxylic acid (I3CA), in the leaf at W3.5 and W5.5 and the MSA at W5.5. Error bars indicate the standard error of the mean of 9-10 biological replicates of leaf samples and four biological replicates of MSA samples. ND means not detected. Statistical groups were assigned using ANOVA followed by Tukey’s post-hoc test for each phytohormone in all genotypes and conditions per stage. Different letters above the barplots indicate significant differences between groups (*p* < 0.05). *RBCS*, *RIBULOSE BISPHOSPHATE CARBOXYLASE SMALL CHAIN*; CS, *CITRATE SYNTHASE*. *IDH*, *ISOCITRATE DEHYDROGENASE*; *FBPase*, *FRUCTOSE-BISPHOSPHATASE*; *PGK*, *PHOSPHOGLYCERATE KINASE*; *RPIA*, *RIBOSE 5-PHOSPHATE ISOMERASE A*; *GADPH*, *GLYCERALDEHYDE 3-PHOSPHATE DEHYDROGENASE*; *PSI*, *PHOTOSYSTEM I*; *PSII*, *PHOTOSYSTEM II;* I3CA, indole-3-carboxylic acid. For the gene filtering and statistics, see Supplementary Datasets 2 and 3.

To corroborate the function of *PPD-H1* in stress resistance, we analysed the levels of the stress-related phytohormones, abscisic acid (ABA) and salicylic acid (SA), and the defence-related indolic compound, indole-3-carboxylic acid (I3CA), at the stages W3.5 and W5.5 in the leaf and at W5.5 in the MSA. HT upregulated the levels of *cis-* and *trans*-ABA and I3CA, in both leaf and MSA of GP, but not in GP-fast (Fig. 4C, D). Similarly, HT increased SA levels in the leaf of GP, but not in GP-fast, while SA levels in the MSA were not affected by genotype or treatment (Fig. 4C, D). These results corroborate the role of *PPD-H1* in stress resistance and trait canalisation in response to temperature variation.

### Genes Involved in Energy Metabolism and Pollen Development Correlate with High Ambient Temperature*-* and *PPD-H1*-Controlled Spike Development and Fertility

Since HT had *PPD-H1*-dependent effects on developmental timing, we searched the leaf and MSA transcriptomes for temperature-regulated developmental genes. HT downregulated transcript levels of *FT1* and its downstream *MADS-BOX* gene *VERNALIZATION 1* (*VRN1*) in the leaf, and *FT2*, the A-class *MADS-BOX* genes *BM3* and *BM8*, and the E-class *MADS34* and *MADS5* in the MSA in all genotypes, while transcript levels were generally higher in wild-type than in mutant genotypes under both temperatures (Supplementary Fig. S11). The expression patterns of these known flowering time regulators did thus not correlate with the HT-induced developmental delays in mutant *ppd-h1* plants versus the accelerated development in wild-type *Ppd-H1* plants. A weighted gene co-expression network analysis (WGCNA) of the MSA DEGs revealed five clusters (C1-C5) with significant correlations to the HT and *PPD-H1*-dependent developmental timing, inflorescence development, and spike fertility (Fig. 5A, Supplementary Fig. S12B, Supplementary Dataset 6). In C3, we identified known stem cell maintenance genes downregulated over development, i.e., *CENTRORADIALIS* (*CEN*), an *ALOG* transcription factor, and *SCARECROW* and *WOX* homologs, which were more strongly downregulated by HT in mutant *ppd-h1* compared to wild-type genotypes (Fig.5A, B, Supplementary Fig. S13A). Similarly, genes in C4, highly expressed over all stages, were more strongly downregulated in the mutant than wild-type genotypes under HT, particularly during floral development (Fig. 5B). This cluster was enriched for transcripts with functions in chromatin assembly and cell cycle and division and was positively correlated with SM induction rates (Fig. 5B, Supplementary Fig. S13B, Supplementary Dataset 5). Interestingly, we found that genes in C2 correlated with the *PPD-H1*-dependent acceleration or delay of inflorescence development under HT (Fig. 5A, B). This cluster was enriched for genes with functions in carbohydrate transport and energy metabolism, i.e., TCA cycle, photosynthesis, glycolysis, and pentose phosphate pathway, which were significantly downregulated under HT in the MSA of mutant *ppd-h1* genotypes but upregulated in the wild-type *Ppd-H1* genotypes (Supplementary Fig. S13C, Supplementary Dataset 5). It included genes encoding for PROTOCHLOROPHYLLIDE OXIDOREDUCTASE A, PHOTOSYSTEM I and II REACTION CENTER SUBUNIT, CHLOROPHYLL A-B BINDING PROTEIN (CAB), which control chlorophyll biosynthesis, light uptake, and energy flow between photosystems, several SWEET sugar transporters, ACID BETA-FRUCTOFURANOSIDASES, which hydrolyzes sucrose into glucose and fructose for sink utilization (Fig. 5B, C, Supplementary Fig. S13C). Furthermore, in accordance with anther and pollen developmental defects observed in mutant *ppd-h1* genotypes under HT, we found anther and pollen development-related genes, i.e., *bHLH89* transcription factor, pollen tube growth regulators *3-KETOACYL-COA SYNTHASE 5* and *PHOSPHOETHANOLAMINE N-METHYLTRANSFERASE 1*, and pollen viability regulator *CALLOSE SYNTHASE 5*, were strongly downregulated in mutant *ppd-h1* genotypes but not in wild-type genotypes at W6.0 under HT (Fig. 5C, Supplementary Fig. S13C). Notably, a number of genes encoding HSPs with possible functions in developmental priming and pollen development (Mareri and Cai, 2022) were specifically expressed at W6.0 and downregulated in the mutant but upregulated in the wild-type genotypes (Supplementary Fig. S13C, Supplementary Dataset 5). Finally, genes with functions in hormone signalling and synthesis, such as *SAUR-LIKE AUXIN-RESPONSIVE PROTEIN*, *GATA* transcription factor, and *YUCCA9* homolog, were downregulated by HT in GP but not in GP-fast (Fig. 5B, C, Supplementary Fig. S13C). These results suggested that HT affected carbohydrate metabolism, hormone signalling, and pollen and anther development in a *PPD-H1*-dependent manner. To corroborate our results, we analysed phytohormone levels of auxin, cytokinins, and gibberellins in the leaf and MSA at W5.5 (Fig. 5D-F, Supplementary Fig. S14). In the MSA, HT reduced indole-3-acetic acid (IAA) levels in both genotypes, with a more pronounced decrease in GP compared to GP-fast (Fig. 5D). Similarly, the levels of major gibberellins, GA44, GA20, and GA8, detected in the inflorescence, were strongly reduced in GP under HT, while in GP-fast, these levels remained comparable to CT samples of GP (Fig. 5E). Notably, GA19 levels increased under HT in GP-fast but were unchanged in GP (Fig. 5E). Additionally, the cytokinin iPR (N6-isopentenyladenosine) was lower in the inflorescence of GP than GP-fast under CT and was further downregulated by HT in GP, but not significantly altered in GP-fast (Fig. 5F). By contrast, the inactive storage form of cytokinin, cZOG (*cis*-zeatin-O-glucoside), increased under HT in GP, but was not significantly affected by HT in GP-fast (Fig. 5F). These results demonstrate that *PPD-H1* controls auxin, gibberellin, and cytokinin levels in the inflorescence under HT to preserve spike fertility and overall reproductive success.

**Figure 5.**
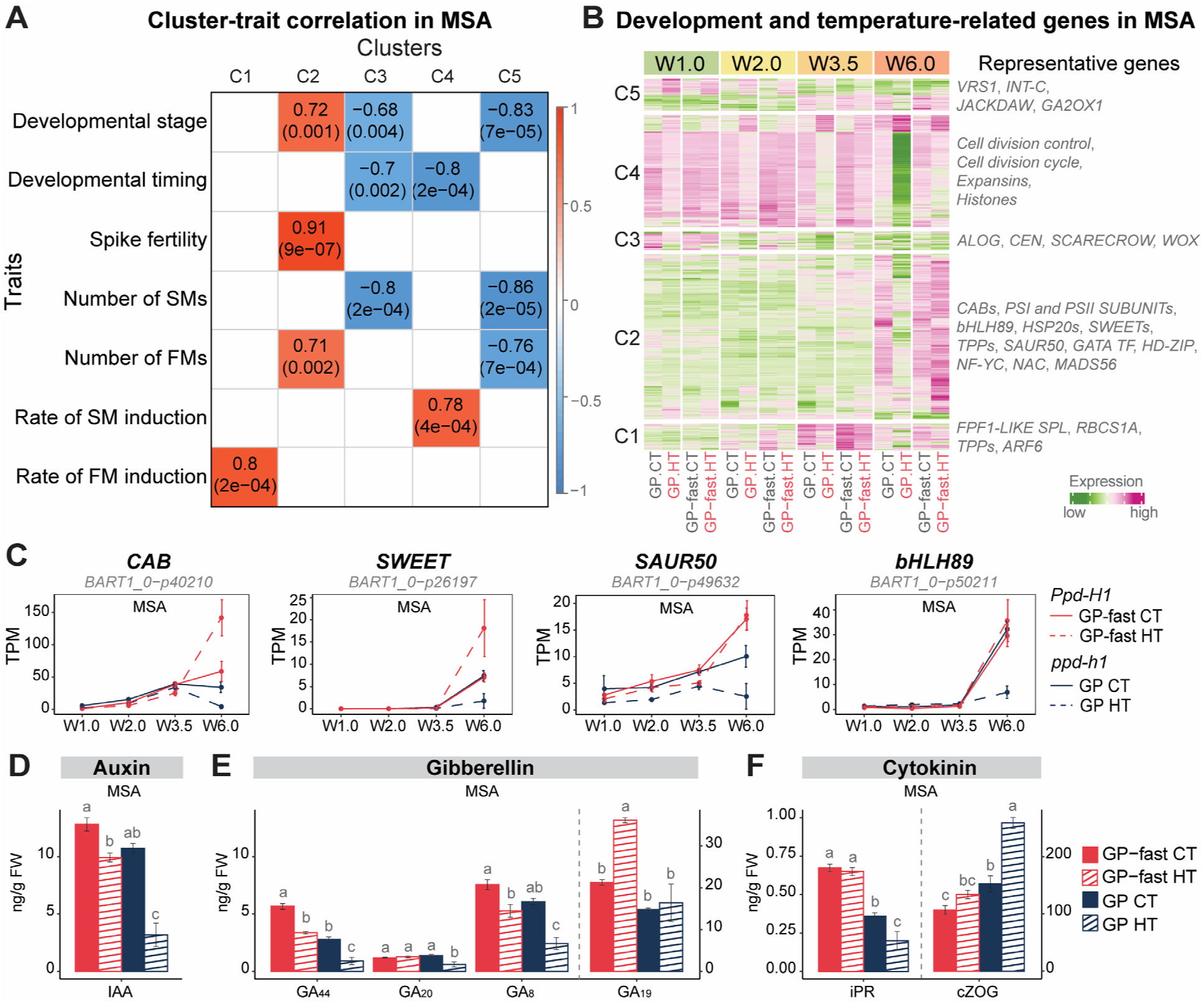
Temperature-Responsive and Development-Related Gene Identification and Phytohormone Changes in the Shoot Apex. (A) Correlation between temperature-responsive gene cluster (column) in the main shoot apex (MSA) and phenotypic trait (row) in spring barley Golden Promise (GP, *ppd-h1*) and its near-isogenic line GP-fast (*Ppd-H1*) grown under control (CT, 20 °C/16 °C, day/night) and high ambient temperatures (HT, 28 °C/24 °C, day/night). Each cell represents the correlation coefficient between a module eigengene (a representative gene expression profile of a module) and a specific trait, with the corresponding p-value provided in parenthesis. The colour scale on the right indicates the strength and direction of the correlation. Positive correlations are shown in red hues, while negative correlations are in blue hues. The intensity of the colour reflects the magnitude of the correlation coefficient, with darker shades representing stronger correlations. (B) Heatmap displays the transcript profiles of genes in Cluster 1 to 5 (C1 to C5) in the MSA of GP (*ppd-h1*) and GP-fast (*Ppd-H1*) under CT and HT. The colour scale represents the range of Z-score normalized mean transcript per million (TPM) values of four biological replicates. The darker green hues represent lower expression, while darker magenta hues represent higher expression relative to other samples in the same gene. The representative gene families are labelled on the side of the heatmap. (C) Transcript profiles of representative genes in C2 that were closely related to developmental timing, spike fertility, and floral meristem (FM) number in the MSA of GP-fast (*Ppd-H1*) and GP (*ppd-h1*) under different ambient temperatures. These genes were involved in light reactions of photosynthesis, sugar transport, auxin signalling, and anther development. Transcript levels are shown in TPM. Solid and dashed lines indicate CT and HT, respectively. Error bars indicate the standard deviation of the mean of four biological replicates. (D-F) Levels of developmental-related phytohormones indole-3-acetic acid (IAA) (D), gibberellins (GAs) (E), and cytokinins (F) in the MSA of GP-fast and GP at W5.5 under CT and HT. Error bars indicate the standard error of the mean of four biological replicates. Statistical groups were assigned using ANOVA followed by Tukey’s post-hoc test for each hormone component. Different letters above the barplots indicate significant differences within the group (*p* < 0.05). SM, spikelet meristem; FM, floral meristem; *CAB*, *CHLOROPHYLL A-B BINDING*; *SWEET*, *SUGARS WILL EVENTUALLY BE EXPORTED TRANSPORTER*; *SAUR50*, *SMALL AUXIN UP-REGULATED RNA 50*; *bHLH89*, *BASIC HELIX-LOOP-HELIX 89;* iPR, N6-isopentenyladenosine; cZOG, *cis*-zeatin-O-glucoside. For the genes associated with development and traits, see Supplementary Datasets 5 and 6.

In summary, the *PPD-H1*-dependent regulation of MSA development by HT correlated with genes involved in stress response, carbon assimilation and metabolism, anther and pollen development, as well as auxin, cytokinin, and gibberellin levels. This suggests that HT affects flowering time by modifying stress resistance and energy metabolism rather than the canonical flowering time regulators like *FT1* in the leaf and *FT2* and floral homeotic genes in the MSA.

## Discussion

Numerous studies have demonstrated that *PHOTOPERIOD 1* (*PPD1*) promotes reproductive development and flowering time under long-day (LD) conditions in barley and wheat by upregulating the expression of Arabidopsis *FT* homologs in the leaf (Turner *et al*., 2005; Beales *et al*., 2007; Digel *et al*., 2016). A natural mutant *ppd-h1* variant with reduced *FT1* expression and delayed flowering time was selected in spring barley, presumably as an adaptation to long growing seasons (Turner *et al*., 2005). A mutation in the CCT domain of *Ppd-H1* was proposed to underlie the flowering time variation between the two major functional *PPD-H1* variants (Turner *et al*., 2005). However, other SNPs outside the CCT domain were also suggested as causal for the *PPD-H1*-dependent flowering time differences in barley (Jones *et al*., 2008). Here, we could confirm the critical role of the CCT domain in controlling developmental timing under LDs while not affecting diurnal clock gene oscillations. The *ppd-h1.1* mutant lacking the C-terminal CCT domain and spring cultivar GP with a single amino acid substitution in the CCT domain flowered at the same time under either CT or HT, and both showed a strong delay in flowering time under HT (Fig. 1). This HT-induced delay in reproductive development was also observed in the spring barley cultivar Scarlett, which carries the same amino acid substitution in the CCT domain as in GP (Supplementary Dataset 1) (Ejaz and von Korff, 2017). The CCT domain of *PRR5* is necessary for association with DNA *in vivo* (Nakamichi *et al*., 2012), and the CCT domain of several circadian clock-regulated *PRRs*, including *PRR7*, bind DNA directly *in vitro* (Gendron *et al*., 2012).

The mutant *ppd-h1* genotypes showed strong phenotypic responses to HT, a delay in flowering time, and a reduction in floret fertility, while the wild-type *Ppd-H1* genotypes showed relatively stable numbers of tillers, spikes, florets, and grains under HT compared to CT. Similarly, mutant *ppd-h1* genotypes showed about twice as many DEGs, and the magnitude of their transcriptional change in response to HT was about two times greater compared to wild-type *Ppd-H1* plants (Fig. 3, Supplementary Datasets 2, 3). The *PPD-H1*-dependent differences in phenotypic and transcriptional plasticity under HT suggest that *PPD-H1* controls environmental canalization or robustness (Waddington 1957; Wagner et al. 1997). This environmental robustness in *Ppd-H1* wild-type genotypes was linked to superior performance under HT compared to the *ppd-h1* mutants (Fig. 1). Pleiotropic effects of *PPD-H1* on adaptation to stress-prone environments, like drought and salt, have been reported before and have commonly been explained by the secondary consequences of developmental timing on growth cycle adaptation to seasonal changes (Wiegmann *et al*., 2019; Gol, Haraldsson and von Korff, 2021; Teklemariam *et al*., 2023). However, our transcriptome and hormone analyses demonstrated that *PPD-H1* controls stress resistance already at early stages in the leaf and shoot apices (Fig. 4). *PPD-H1* belongs to the family of *PESUDO RESPONSE REGULATOR* (*PRR*) genes, which regulate circadian rhythms, light signalling and stress response (Nakamichi *et al*., 2009, 2010). In Arabidopsis, *PRR7* was identified as a transcriptional repressor of genes involved in ABA-dependent and independent stress responses (Liu *et al*., 2013). Triple *prr579* mutants showed increased drought, high salinity, oxidative, and cold stress tolerance in different species linked to alterations in circadian timing and, thus, gating of gene expression (Nakamichi *et al*., 2009). However, in barley, the mutant *ppd-h1* genotypes did not show alterations in diurnal and circadian oscillations (Campoli *et al*., 2012) (Supplementary Fig. S2). Furthermore, the upregulation of stress-related genes and ABA levels were linked to higher leaf senescence rates and lower floret fertility in the mutant *ppd-h1* plants (Fig. 2, 4, Supplementary Fig. S4, S7). We thus conclude that mutant *ppd-h1* plants were more stress-susceptible, which resulted in the upregulation of stress response genes and hormones in the leaf and MSA (Fig. 4). This finding contrasts with observations in Arabidopsis, where *prr* mutants were described as more stress tolerant due to the derepression of stress response genes and increased ABA biosynthesis (Nakamichi *et al*., 2009; Liu *et al*., 2013). These differences in the effects of *PRR* genes on stress resistance between species might be due to the differential effects of *prr* mutations on the circadian clock. The *ppd-h1* mutants in barley show no circadian or diurnal misregulation of clock or clock target genes, while Arabidopsis *prr* mutants are impaired in circadian oscillations (Farré *et al*., 2005; Campoli *et al*., 2012; Müller *et al*., 2020). Additionally, differences in the stress application and trait evaluation might explain the differences: short-term lethal stress and survival rates in Arabidopsis, and long-term mild stress and reproductive development in barley (Nakamichi *et al*., 2009; Liu *et al*., 2013).

Our objective was to identify leaf-and MSA-specific developmental genes and molecular networks that correlate with the distinct HT responses of *PPD-H1* variants, which either accelerate or delay flowering time in response to HT. However, HT reduced transcript levels of major developmental genes in both leaves and MSAs, and we did not detect further developmental genes that correlated with the *PPD-H1*-dependent delay or acceleration of development under HT in the global transcriptomes (Supplementary Fig. S11). For example, *FT1* in the leaf, and *FT2* in the MSA were downregulated by HT in all genotypes (Supplementary Fig. S11). This contrasts with Arabidopsis, where HT accelerates flowering under non-inductive short-day conditions through the upregulation of *FT* in leaves (Balasubramanian 2006). Our MSA transcriptome analysis revealed that transcript levels of genes involved in the TCA cycle, Calvin cycle, light reactions of photosynthesis, glycolysis, and pentose phosphate pathway were higher in wild-type *Ppd-H1* plants compared to mutant *ppd-h1* plants not only under HT, but also under CT (Fig. 4). Moreover, HT downregulated the expression these genes in mutant *ppd-h1* plants but upregulated or did not affect their expression in wild-type *Ppd-H1* plants (Fig. 4). These transcriptional changes were strongly associated with developmental differences controlled by *PPD-H1* and ambient temperature interaction, such as developmental timing, SM and FM induction rates, and spike fertility (Fig. 5). These findings suggest that *PPD-H1* controls energy and carbon metabolism, as also seen in the differential expression of carbon signalling and transport genes. Genes encoding CAB, PHOTOSYSTEM I and II SUBUNIT proteins, and SWEET were downregulated by HT in the mutant *ppd-h1* plants but upregulated in the MSA of wild-type *Ppd-H1* plants (Supplementary Dataset 5).

The downregulation of photosynthesis-related *CABs* in the inflorescence has already been linked to floral abortion and anther defects in the barley *tip sterile 2.b* (*tst2.b*) mutant, possibly because the plastidial energy supply to developing floral tissues was affected (Huang *et al*., 2023). This is in line with our observation that HT reduced starch content in the pollen grains, and several sugar invertase genes (e.g., *BETA-FRUCTOFURANOSIDASE*) were downregulated in the MSA of mutant *ppd-h1* plants but not in wild-type plants (Supplementary Fig. S13). However, even before pollen development, a strong upregulation of ABA and downregulation of cytokinin and cell cycle genes in the mutant *ppd-h1* genotypes suggested that cell cycle and, thus, cell proliferation was strongly repressed in the mutant inflorescences under HT, but not in the wild type (Fig. 4, 5, Supplementary Fig. S13). Moreover, *ppd-h1* mutants but not wild-type plants showed a reduction in auxin levels in the inflorescences under HT, which impairs anther development and male sterility in barley (Sakata *et al*., 2010; Oshino *et al*., 2011). These findings suggest that mutant *ppd-h1* genotypes display a strong and comprehensive stress response that changes developmental timing, spike growth and development, energy supply for anther and pollen development in the spike, and thus floret fertility. Interestingly, even under CT conditions, *ppd-h1* mutants displayed reduced fertility and transcriptomic changes, indicative of a stress response compared to the wild-type genotypes (Fig. 2, Supplementary Fig. S13). We speculate that the wild-type *Ppd-H1* allele enhances stress resistance, thereby maintaining high carbon utilization and energy supply to drive rapid reproductive development and mitigate the adverse effects of HT on floret fertility.

The natural mutant *ppd-h1* was selected in spring barley varieties widely grown in Middle and Northern Europe because it delays reproductive development and thereby increases tiller and spike number and the number of spikelets, florets and grains per spike (Turner *et al*., 2005; Digel, Pankin and von Korff, 2015). However, under less favourable conditions, the increased stress susceptibility of these *ppd-h1* natural mutants outbalances the yield advance under favourable conditions. *PPD-H1* is thus an interesting target to dissect further the interrelationship between development, energy metabolism, and stress resistance and to breed climate-adapted barley varieties.

## Materials and Methods

### Plant Materials and Growth Conditions

In this study, we selected two genotypes carrying a wild-type *Ppd-H1* allele and three genotypes with mutant *ppd-h1* allele to study the interaction between *Ppd-H1* and HT on shoot growth and inflorescence development. Golden Promise (GP) and Scarlett are spring barley cultivars carrying a natural mutation in the CCT domain of *PPD-H1* (G>T at CDS position 1969), according to gene model *HORVU.MOREX.r3.2HG0107710.1* (Supplementary Dataset 1), causing a Gly-to-Trp amino acid exchange, and resulting in a late flowering time under long-day (LD; 16 h/8 h, day/night) conditions. GP-fast and S42-IL107 are two near-isogenic lines (NILs) derived from GP and Scarlett, respectively, carrying wild-type *Ppd-H1* alleles from winter barley Igri and wild barley (*Hordeum vulgare* ssp. *spontaneum*) accession ISR42-8 (Supplementary Dataset 1). These lines are early flowering under LD (Schmalenbach *et al*., 2011; Digel, Pankin and von Korff, 2015; Gol, Haraldsson and Von Korff, 2021).

Plants were sown in a 96-well growing tray with ED73 soil (Einheitserde Werkverband e.V., Germany) mixed with 7% sand and 4 g/L Osmocote Exact Hi.End 3-4M (ICL Group Ltd.). Seeds were stratified for three days at 4 °C in the soil for even germination and then transferred to controlled growth chambers (PAR 300 µM/m^2^s, humidity 60%) with inductive LD conditions (16 h/8 h, day/night) set to control (CT; 20 °C/16 °C, day/night) or high ambient temperature (HT; 28 °C/24 °C, day/night).

### Generation of an induced *PPD-H1* mutant using CRISPR/Cas9

To extend our study beyond natural *PPD-H1* alleles, we used CRISPR/Cas9 to introduce new mutations. RGEN Tools Cas-Designer (Bae, Park and Kim, 2014; Park, Bae and Kim, 2015) was used to select two single guide RNAs (sgRNAs) close to or within the conserved CCT domain of *PPD-H1* (Figure S1). The sgRNAs were cloned into transformation vectors according to Kumar et al. (2018) and Helmsorig et al. (2023). GP-fast embryos carrying the wild-type *Ppd-H1* allele were used for transformation, which was performed according to the protocol of Hensel et al. (2009). M0 plants were screened by PCR for the presence of Cas9 by primer sequences 5’-GAGCGCATGAAGAGGATCGA-3’ and 5’-GGACACGAGCTTGGACTTGA-3’. and genotyped for mutations using *PPD-H1*-specific primer sequences 5’-TGTGTGCGGTCTCCAATCAA-3’ and 5’-GCACCTGCAAAACGAACGAC-3’ covering the sgRNA-targeted region through Sanger sequencing. One line carrying a homozygous 1-bp T-insertion at CDS position 1371 (according to the gene model *HORVU.MOREX.r3.2HG0107710.1*) in the M0 was selected for further analysis and hereafter named *ppd-h1.1*. The T-insertion leads to a frameshift and inserts an immediate premature stop codon, resulting in a truncated protein size of 444 amino acids. The M1 generation was used to remove the transformation vector and *ppd-h1.1* plants in the M2 generation were analysed in this study.

### Comparison of *PPD-H1* alleles

RNAseq reads obtained from leaf data at W3.5 in CT and HT were mapped jointly per genotype to the Morex v3 reference (Mascher *et al*., 2021) using the STAR aligner (version 2.7.10b) (Dobin *et al*., 2013). The resulting BAM files were indexed with SamTools (version 1.18) (Li *et al*., 2009). Examination of polymorphisms within the *PPD-H1* gene was performed using the Integrative Genomics Viewer (IGV, version 2.17.4) (Robinson *et al*., 2011). Polymorphisms were extracted based on the gene model *HORVU.MOREX.r3.2HG0107710.1*, which also were used to verify the 1-bp insertion in the induced *ppd-h1.1* mutant line. The resulting CDS sequences and translated protein sequences of the *PPD-H1* alleles were compared by multiple sequence analysis using CLUSTAL Omega (version 1.2.4) (Madeira *et al*., 2022).

### Diurnal gene expression analysis

We examined the effect of the mutation in *Ppd-H1* on the circadian oscillations in GP-fast, GP, and *ppd-h1.1*. Plants were grown under LD conditions until 14 days after emergence (DAE). Plants were sampled every 4 h over a complete light/dark cycle of 24 h (light from ZT0-16, dark from ZT16-24). For each replicate, the middle sections of the youngest, fully elongated leaf of two plants were pooled. Three biological replicates were sampled.

RNA was extracted by grinding the samples using TRIzol (Thermo Fisher Scientific) according to the manufacturer’s instructions. RNA was resuspended in 60 µL of diethyl dicarbonate-treated water at 4 °C overnight. The remaining DNA was removed by subsequent DNAse I treatment (Thermo Fisher Scientific) according to the manufacturer’s instructions. cDNA was synthesized on 2 µg of total RNA using ProtoScript II First Strand cDNA Synthesis Kit (NEB) following the manufacturer’s instructions. Gene expression levels were determined by RT-qPCR in a LightCycler 480 (Roche) using gene-specific primers (Supplemental Table S2). The reaction was performed using 4 µL of cDNA, 5 µL of 2X Luna qPCR Master Mix (NEB), 0.02 mM of forward and reverse primer, and 0.75 µL of water with the amplification conditions 95 °C for 5 min, 45 cycles of 95 °C (10 s), 60 °C (10 s) and 72 °C (10 s). Non-template controls were added to each plate, and dissociation analysis was performed at the end of each run to confirm the specificity of the reaction.

Two technical replicates were used and averaged in analyses for each biological replicate. The expression of *HvACTIN* was used as a reference to calculate the relative gene expression of the target genes.

### Phenotyping

To score the development of the main shoot apex (MSA) under CT and HT, five representative plants per genotype and treatment were dissected every 3 to 4 days from emergence to post-pollination. The developmental stage of the MSA was scored according to the Waddington scale (Waddington et al. 1983). This scale rates the progression of spikelet initiation and then the development of the most advanced floral primordium and the pistil of the MSA.

Flowering time was scored in DAE until the awns emerged from the flag leaf, called awn tipping (Zadoks, Changt and Laboratory, 1974). The shoot phenotypes, such as plant height and the number of tillers and spikes were measured on 15-20 plants for GP and GP-fast, and on 11-15 plants for Scarlett and S42-IL107 in each condition at plant maturity. The spike phenotypes, such as the number of florets and grains, were measured on the MSA from the same plant scored for the shoot phenotypes. Spike fertility was determined by the ratio of grain number to the number of final florets. The length, width, and thousand-grain weight (TGW) were measured using seeds from three individual plants per genotype and condition using the MARVIN Seed Analyser (GTA Sensorik, Germany).

To score the rate of spikelet meristem (SM) and floral meristem (FM) induction under LD, 16-23 MSAs of GP and GP-fast, 3 MSAs from *ppd-h1.1*, and 8-17 MSAs of Scarlett and S42-IL107 under CT and HT were dissected at the spikelet induction stage (W2.0), the stamen primordium stage (W3.5), the pistil primordium stage (W4.5), and the style primordium stage (W6.0). SMs were scored as soon as the spikelet induction stage (W2.0) began. The number of SMs included those that had differentiated into floral primordia or floral organs. An FM was scored as soon as the stamen primordium was initiated (W3.5), and the number of FMs included those that had differentiated floral organs or developed into florets.

To score the leaf appearance rate (LAR), the number of emerged leaves on the main culm was recorded every 2-3 days. LAR was calculated as the time interval between the sequential emergence of leaves on the main stem on 10 plants per genotype and treatment. The length and width of all leaf blades on the main culm were measured when they were fully elongated, as seen when the auricle of the leaf was widely open. Leaves from 10 individual plants per genotype and treatment were measured.

To score the senescence rate of the leaf, we collected the second leaf (L2) of GP-fast, GP, and *ppd-h1.1* grown under CT and HT at 19, 27, and 32 DAE. 10-16 plants were scored per genotype and temperature combination. Based on the percentage of the yellow or brown area in the leaf, we defined the score for the senescence as 1 (area ≤ 1/3), 2 (1/3< area ≤ 2/3), and 3 (area > 2/3).

To analyze the cell size of the leaf blade, epidermal imprints were taken of the fully elonged L2 from three plants per genotype and condition. To minimize the positional effect on the cell size, we selected two adaxial positions at 1/3 from the leaf tip and the base and brushed an area length of ca. 2 cm with nail polish. After drying, epidermal imprints were removed carefully with transparent tape and attached to microscope slides. Images of the imprints were captured, and 30 lateral cells (Wenzel et al., 1997) at each position were measured for the cell length. Cell width was calculated by the number of cells within 600 pixels (0.822 pixel/µm) horizontal distance using the software Fiji (Schindelin et al., 2012).

To examine the effects of HT on anther development, anther length was measured in the MSA at W9.0 from three serial florets at the top, middle, and basal position. The average of three serial florets represents the anther size at each position per biological replicate. Anthers from three plants per genotype and condition were scored. The length of anthers was measured using Fiji (Schindelin *et al*., 2012).

To measure the pollen viability, stamens from the central floret located at the middle of MSA were collected at W10.0 from five plants of GP and GP-fast and from three plants of Scarlett and S42-IL107. The Alexander’s staining solution consists of 10 mL 95% Ethanol, 1 mL 1% malachite green, 55.5 mL distilled water, 25 mL glycerol, 5 mL 1% acid fuchsin, 0.5 mL 1% orange G, 4 mL glacial acetic acid, and stored in the dark (Peterson, Slovin and Chen, 2010). Stamens were collected into a 1.5 mL tube and 50 µL of Alexander’s Staining solution was added, followed by incubation at 100 °C for 40 min before imaging. Anthers were cut into 3 pieces on a microscope slide with a sharp scalpel and collected into a 1.5 mL tube with 50 µL of Alexander’s Staining solution. Pollen was released by vortexing for 10 s, and then 10 µL of the polle-staining mixture was taken onto the microscope slide for imaging. The number of pollen was counted using the ‘Analyze Particles’ function in Fiji (Schindelin *et al*., 2012).

To document the development of the MSA, the length of anthers, and the number of viable and inviable pollens, we used a Nikon DS-Fi2 digital camera attached to a Nikon SMZ18 stereo microscope to capture the images.

### Inflorescence Meristem Size Measurement

To compare the inflorescence meristem (IM) width and height, three to four MSAs from GP and GP-fast grown under CT and HT were collected from the spikelet induction stage (W2.0) until the stamen primordium stage (W3.5) every 2-4 days. To process the MSAs for the 3D reconstruction by confocal microscopic imaging, MSAs were fixed in 2 mL ice-cold 4% paraformaldehyde in 1% phosphate buffered saline (PBS), and vacuum was applied for 15 min two times on ice. Then the fixative was replaced with a fresh 2 mL 4% para-formaldehyde in 1% PBS, and the MSAs were kept in the fresh fixative at 4 °C overnight. The next day, the MSAs were washed in 1% PBS four times to remove the fixative and then treated with ClearSee solution (Kurihara *et al*., 2015) for seven days. The treated MSAs were stained with 0.1% SR2200 cell wall stain (SCRI Renaissance 2200) (Musielak *et al*., 2015), before imaging using a confocal laser scanning microscope (Olympus, FV3000). The stained MSAs were imaged using a 30X silicon immersion objective, excited at 405 nm, and detected between 415 and 476 nm. The IM border was defined by the first cell in the outermost cortex cell layer that doubled in cell length compared to its distal neighbor (Kirschner *et al*., 2018) using Fiji Software (Schindelin *et al*., 2012).

### Phytohormone extraction

We analysed the phytohormone levels in GP-fast and GP plants grown under CT and HT. Plants were grown in 7*7*7 cm^3^ pots. For the leaf tissue, the entire latest and fully elongated leaf blade on the main culm and 5 mm of adjacent leaf sheath were collected at W3.5 and W5.5, respectively. Four plants were pooled together for each biological replicate. 9-10 biological replicates were examined. For the inflorescence tissue, 8 MSAs were collected at W5.5 and pooled together for each biological replicate. Four biological replicates were used for the measurement.

All samples were collected, immediately frozen, and ground into fine powder with a mortar and pestle using liquid nitrogen. For each sample, 40 to 50 mg of ground tissue was used for the extraction. The extraction procedure was based on Pan et al. (2010), with optimizations regarding alterations in the internal standard mix composition, extraction with an ultrasonic bath, and treatment of the dried phytohormone extracts for optimized LC-separation.

100 µL extraction solution 1 (isopropanol/H_2_O/37%(w/w) HCl, 2:1:0.002; v/v/v), including the internal standard mix, and 400 µL extraction solution 2 (isopropanol/H_2_O/37 %(w/w) HCl, 2:1:0.002; v/v/v) without the internal standard were added to the ground plant material, followed by immediate vortexing. The extraction procedure was carried out in an ice-cooled ultrasonic bath for 30 min (35 kHz, Power 160 W) while vortexing every 10 mins. Then 1 mL methylene chloride was added and the sonication step was repeated as described above. Phase separation was achieved by centrifugation for 5 min at 21000 x g and 4 °C. The lower phase was transferred into a new 2 mL sample tube and concentrated using a Techne™ sample concentrator with nitrogen flow. The dried extracts were redissolved immediately in 50 µL methanol. 50 µL of H_2_O was added to improve LC-separation of those phytohormones eluting at early retention times. The extracts were kept at 4 °C overnight and were subsequently centrifuged for 5 min (21000 x g, 4 °C) to remove precipitated compounds. The clear supernatant (approx. 90 µL) was used for LC-MS analysis of phytohormones. Samples were analysed directly after extraction or stored at -20 °C until analysis.

### LC-MS Analysis of Phytohormones

10 µL sample volumes of phytohormone extracts were used for injection. Phytohormones were separated on a Dionex HPG 3200 HPLC system (Thermo Scientific, Germany) with a binary gradient system.

LC-separation of cytokinins was done on a C18 Atlantis™ Premier BEH column, 2.5 µm, 2.1 mm X 150 mm (Waters, Ireland). All other phytohormones were separated on a C18 XSelect™ HSS T3 Column, 2.5 µm, 3 mm X 150 mm (Waters, Ireland).

The binary gradient system for both HPLC-columns was as follows: mobile phase A consisted of water and 0.1% formic acid (FA), and mobile phase B consisted of methanol and 0.1% FA. The flow rate was 0.35 mL/min. Starting conditions were 2% mobile phase B, increased to 99 % B from 1-18 min. The plateau was held for 4 min and the gradient was returned to 2% mobile phase B within 1 min. The system was equilibrated at 2% B for 7 min prior to the next injection.

Phytohormones were analyzed with a maXis 4G Q-TOF mass spectrometer (Bruker Daltonics, Germany) equipped with an ESI source. The operating conditions were as follows: dry gas (nitrogen): 8.0 L/min, dry heater: 200 °C, nebulizer pressure: 1.0 bar, capillary voltage: 4500 V. Data was acquired in stepping mode with collision Rf voltage ranging from 300 to 500 Vpp for optimal signal intensities of compounds between 100-500 m/z.

Cytokinins were analyzed in positive ion mode; salicylic (SA), abscisic acid (ABA), indole-3-carboxylic acid (I3CA) and gibberellins (GAs) were analysed in negative ion mode. The auxin indole-3-acetic acid (IAA) was measured in both negative and positive ion modes, but showed a better signal-to-noise ratio in the positive ion mode, which was therefore used for IAA quantification.

### Identification and Quantification of Phytohormones

Phytohormones were identified by their retention time (RT) compared with the RT of reference standards and the measured m/z compared with the calculated m/z. Standards were purchased from OlChemim Ltd. (Czech Republic) except for SA and IAA which were purchased from Sigma-Aldrich (Germany) and Supelco (Germany). Calibration curves were measured from 1 nM to 750 nM for all phytohormone standards. Isotopically labelled standards from OlChemim Ltd. (Czech Republic) were used as internal reference standards at a fixed final concentration of 75 nM. The same internal standard mix, containing 0.075 nmol per 100 µL of each labelled standard, was used during sample extraction and for the calibration curves. Absolute quantification was achieved by using the slopes of calibration curves plotted from area ratios of the unlabelled standards and their respective isotopically labelled standards (Supplementary Table S3). Where no corresponding isotopically labelled standard was available, a structurally related standard was used: d_2_-GA19 was used for GA8 and d_5_-tZOG for cZOG. (±)-*cis*,*trans-*ABA was used as a standard for calibration curves to quantify the natural isomer (+)-*cis*,*trans*-ABA. Peak areas were determined with the Software QuantAnalysis (version 5.3, Bruker Daltonics, Germany) and further calculations were done in Microsoft Excel 2010.

### Statistical Test and Plotting of the Phenotypic Traits

An ANOVA test followed by a Tukey’s HSD test was conducted to compare differences among genotype and condition groups using the ‘Agricolae’ R package (Mendiburu and Team, 2017). A Student’s *t*-test was conducted to compare the traits between different temperatures within the same genotype. The distribution of data points was presented by boxplots using ‘ggplot2’ R package (Wickham, 2009).The box was bound by the lower (Q1) and the upper (Q3) quartiles, and the length of the box represents the interquartile range (IQR). The lower, middle, and upper quartiles were the values under which 25%, 50%, and 75% of data points were found when they were arranged in increasing order. The whisker of the boxplot was calculated by Q1-1.5*IQR (lower) or Q3+1.5*IQR (upper). The values outside the whiskers were plotted separately as dots and considered suspected outliers. The development of the MSA, the rates of leaf appearance, SM and FM induction were presented by regression lines using the function *geom_smooth()* of the ‘ggplot2’ R package (Wickham, 2009) in combination with the argument *method = lm*.

### Whole Transcriptome Sequencing

To detect changes in the global leaf and MSA transcriptomes in response to HT, we harvested at ZT14 the leaves and developing MSAs of GP and GP-fast grown under LD either under CT or HT at four developmental stages, W1.0, W2.0, W3.5, and W6.0 for RNA-sequencing. Each replicate of leaf samples was collected by pooling the latest fully elongated leaf on the main culm from three plants. Each replicate of MSA samples was collected by pooling ca. 30, 20, 15, and 10 MSAs from plants at stages W1.0, W2.0, W3.5, and W6.0, respectively. All MSA samples were collected under a stereo microscope in the environment where the plant grew. A total of three to four biological replicates of MSA and leaf samples were used for RNA-sequencing.

Total RNA was extracted from MSA and leaf samples using the RNeasy Plant Mini Kit (Qiagen, Germany) with beta-mercaptoethanol added, following the manufacturer’s instructions, and followed by the digestion with DNase I (Qiagen, Germany). RNA samples passing a cutoff of RNA Integrity Number (RIN) ≥ 6 were used for mRNA library preparation with poly(A)-enrichment. Paired-end 150 bp sequencing was performed on NovaSeq 6000 sequencing platform, and at least 5G of clean reads data per sample were generated by Novogene Co., Ltd (UK). All samples passed the assessment of the adaptor and GC contents by FastQC (Andrews 2010).

### Reads Mapping and Gene Annotation

To quantify transcripts, all cleaned reads were mapped to the BaRT1 reference (Rapazote-Flores *et al*., 2019) using Salmon (v. 0.14.1) (Patro *et al*., 2017), and the mapping rates were estimated with an average of ∼93%. We kept transcripts with a minimum of 1 CPM (counts per million) in at least three samples. Normalization of TPM (transcript per million) was conducted using the ‘ThreeDRNAseq’ R package (v2.0.1) (Guo *et al*., 2021). For TPM values, see Supplementary Dataset 2 and 3. As the samples were collected at different DAEs and RNA extraction was conducted in several batches, we removed the batch effect in the gene count by using the ThreeDRNAseq’ R package (v2.0.1) (Guo *et al*., 2021).

To improve the annotation of the reference, we aligned the CDS sequences of BaRT1 with BLASTX (v2.12.0) against the peptide sequences of *Arabidopsis thaliana* TAIR10 (Cheng *et al*., 2017), with pident of 25 and *e*-value of 10^-6^. A reciprocal BLASTN (v2.13.0) (Camacho *et al*., 2009) alignment was conducted using the longest transcript CDS sequence per gene of barley BaRT1 (Rapazote-Flores *et al*., 2019) and Morex v3 (Mascher *et al*., 2021). A best hit was selected by e-value and bitscore, and a reciprocal match was identified when there was a best hit between gene models in both directions. For the gene annotation and identifiers in other references, see Supplementary Dataset 2 and 3.

### Identification of Temperature-Responsive Genes in the Leaf

To identify differentially expressed genes (DEGs) in response to HT in the leaf, a three-way ANOVA was calculated using TPMs from leaf samples of GP and GP-fast. Genes with a Benjamini-Hochberg (BH) procedure adjusted *p*-value below 0.01 for the temperature or temperature* genotype, or temperature*genotype*stage effects were then subjected to pairwise comparisons (HT versus CT) for detecting significant transcriptional changes between HT and CT within each genotype and developmental stage, using the count-based Fisher’s Exact Test in the R package ‘EdgeR’ (v3.32.1) (Robinson, McCarthy and Smyth, 2010). The FDR of the comparison was adjusted by the BH procedure, thus the genes with BH.FDR < 0.01 and |log_2_FC| ≥ 1 in at least three developmental stages were referred to as temperature-responsive genes. To confirm the thermoresponsiveness of the genes detected in GP and GP-fast, we tested the transcriptional changes of these genes in *ppd-h1.1*, Scarlett, and S42-IL107 at W3.5 and W6.0, respectively. Thus, genes with BH.FDR < 0.01 and |log_2_FC| >0 at either W3.5 or W6.0 in *ppd-h1.1*, Scarlett, and S42-IL107 were further analysed. To reduce the interference from genes other than *PPD-H1*, we focused on genes exhibiting up-or down-regulation in response to HT shared between GP and *ppd-h1.1* or Scarlett, and between GP-fast and S42-IL107. For detailed information on statistics and results, see Supplementary Dataset 2.

### Identification of Temperature-Responsive Genes in the MSA

To identify DEGs in response to HT in MSA samples, a three-way ANOVA followed by the count-based Fisher’s Exact Test was conducted in the MSA samples of GP and GP-fast, as described above for the leaf samples. Genes with BH.FDR <0.01 and |log2FC| ≥ 1 were referred to as temperature-responsive genes. We analysed the temperature-responsive genes separately by different developmental stages. To confirm the thermoresponsiveness of the genes detected in GP and GP-fast, we tested the significance of the transcriptional differences of the genes in the MSA of *ppd-h1.1*, Scarlett, and S42-IL107 at W3.5 and W6.0, respectively, using the pairwise comparison (HT versus CT). Thus, genes with BH.FDR < 0.01 and |log_2_FC| > 0 at either W3.5 or W6.0 in *ppd-h1.1*, Scarlett, and S42-IL107 were further analysed. To reduce the interference from genes other than *PPD-H1*, the temperature-responsive genes shared between (a) the wild-type *Ppd-H1* genotypes (GP-fast and S42-IL107) at W3.5 and W6.0, and (b) mutant *ppd-h1* genotypes (GP, *ppd-h1.1*, Scarlett) at W3.5 and W6.0, and the gene in GP and GPfast at W1.0 and W2.0. For detailed information on statistics and results, see Supplementary Dataset 3.

### Identification of *PPD-H1*-Dependent Genes in the Leaf

To identify *PPD-H1*-dependent DEGs in the leaf, a three-way ANOVA was calculated using TPMs for leaf samples of GP and GP-fast, as described above. Genes with a BH procedure adjusted *p*-value below 0.01 for the genotype, or genotype*temperature, or genotype*temperature*stage effects were then subjected to pairwise comparisons (GP-fast versus GP) for detecting significant transcriptional changes between GP-fast and GP at each developmental stage under CT and HT conditions, respectively, using the count-based Fisher’s Exact Test in the R package ‘EdgeR’ (v3.32.1) (Robinson, McCarthy and Smyth, 2010). The FDR of the comparison was adjusted by the BH procedure, thus genes with BH.FDR < 0.01 and |log_2_FC| ≥ 1 in at least three developmental stages under either CT or HT were referred to as a *PPD-H1*-dependent genes. To confirm the *PPD-H1*-dependent regulation of the gene expression, we tested their transcriptional changes in GP-fast versus *ppd-h1.1* and in S42-IL107 versus Scarlett. Thus, genes with BH.FDR < 0.01 and |log_2_FC| >0 at either W3.5 or W6.0 in *ppd-h1.1*, Scarlett, and S42-IL107 were further analysed. To reduce the interference from the genes other than *PPD-H1*, we focused on genes exhibiting the same regulation shared (a) between GP-fast versus GP and S42-IL107 versus Scarlett or (b) between GP-fast versus GP and GP-fast versus *ppd-h1.1*. For detailed information on statistics and results, see Supplementary Dataset 2.

### Identification of *PPD-H1*-Dependent Genes in the MSA

To identify *PPD-H1*-dependent DEGs in the MSA, a three-way ANOVA followed by the count-based Fisher’s Exact Test was conducted in the MSA samples of GP and GP-fast, as described above for the leaf samples. The genes with BH.FDR <0.01 and |log2FC| ≥ 1 were referred to as *PPD-H1*-dependent genes. We analysed the *PPD-H1*-dependent genes separately by temperature. To confirm the *PPD-H1*-dependent regulation in gene expression, we conducted pair-wise comparisons in GP-fast versus *ppd-h1.1* and in S42-IL107 versus Scarlett. Genes with a significant difference in gene expression at BH.FDR < 0.01 and |log_2_FC| > 0 were further analysed. To reduce the interference from genes other than *PPD-H1*, we focused on the genes shared across (a) the wild-type *Ppd-H1* genotypes (GP-fast and S42-IL107) at W3.5 and W6.0, and (b) mutant *ppd-h1* genotypes (GP, *ppd-h1.1*, and Scarlett) at W3.5 and W6.0, and (c) the genes in GP and GPfast at W1.0 and W2.0. For detailed information on statistics and results, see Supplementary Dataset 3.

### Co-expression analysis

TPM data of temperature-responsive genes in the MSA sample of GP-fast and GP were used to construct co-expression networks, identify modules, and determine module-trait relationships. Average TPM values of four biological replicates at each developmental stage and temperature condition were used to identify co-expression modules using the WGCNA package in R (Langfelder and Horvath, 2008). Co-expression modules were identified using the TOMtype parameter unsigned. A Pearson correlation matrix was calculated for all pairs of selected genes, and an adjacency matrix was constructed by raising the correlation coefficients to the twelve power. The minModuleSize parameter was set to 30, and modules were identified using the Dynamic Hybrid Tree Cut algorithm with MergeCutHeight at 0.25, which identified nine modules (Supplementary Dataset 6).

### Gene Ontology Enrichment Analysis and Plotting

To identify significantly enriched gene ontology (GO) terms, we conducted GO enrichment analyses using the ‘GOEnrichment’ tool of the Triticeae Gene Tribe database (Chen *et al*., 2020) with BH.*p*-value < 0.05. For the complete information on all GO terms and genes in each GO term, see Supplementary Dataset 4. Heatmaps for GO enrichment were generated with the -log_10_(FDR) values of the top ten biological function GO terms using the ‘ThreeDRNAseq’ R package (v2.0.1) (Guo *et al*., 2021) in combination with ComplexHeatmap (Gu, Eils and Schlesner, 2016). The distance matrix was computed using the Euclidean method. Heatmaps of gene expression in selected GO terms were generated with the log_10_(TPM+1) values using the same packages as described above. For the complete gene list for the heatmaps, see Supplementary Dataset 5. Principle component analysis (PCA) was conducted using the ‘ThreeDRNAseq’ R package (v2.0.1) after removing batch effects (Guo *et al*., 2021).

## Author Contributions

T.L. and M.v.K. conceived and designed the experiments. T.L. conducted experiments, plant phenotypic analyses, RNA-sequencing analyses, and interpreted the data. A.W. designed single-guide RNAs, and G.B. performed transformations. A.W. and G.B. genotyped the mutants. A.W. conducted the diurnal expression experiment and analysed the gene expression. T.L. and A.W. characterised the *ppd-h1.1* mutant and conducted the RNA-sequencing experiment in *ppd-h1.1*. K.K. conducted the phytohormone experiment with the help of V.T. and analysed the data. V.W. and S.M. established the protocol for phytohormone measurement and analysed the LC-MC results. I.V. assisted confocal microscopy and image analysis. E.B.H. improved the gene annotation and analysed RNA-sequencing data. G.H. assisted with sample collection, trial experiment, and experimental setup. R.S. provided guidance, feedback and support throughout the project. T.L. wrote the manuscript with the help of A.W., R.S., and M.v.K.

## Acknowledgements

We greatly thank Thea Rüjes, Rebekka Schüller, and Nina Döring for excellent technical assistance, Maria Bernal, Jacqueline Baum, and Anna Bechler for verifying phenotypes and diurnal expression of induced mutant, the Center for Advanced Imaging (CAi), especially Sebastian Hänsch and Stefanie Weidtkamp-Peters, for their expertise on confocal imaging and analysis, Dominik Brilhaus for data management support, and Jinshun Zhong for critically reading this manuscript.

## Supplementary Data

**Supplementary Figure S1.** Summary of CRISPR/Cas9-Induced Mutation in *Ppd-H1*.

**Supplementary Figure S2.** Effects of *PPD-H1* on Diurnal Gene Expression Pattern of Selected Circadian Clock Genes.

**Supplementary Figure S3.** Effects of High Ambient Temperature on Shoot Growth in Scarlett and S42-IL107.

**Supplementary Figure S4.** Effects of High Ambient Temperature on Leaf Development and Senescence.

**Supplementary Figure S5.** Effects of High Ambient Temperature and *PPD-H1* on the Spikelet Meristem and Floral Meristem Induction Rate in Scarlett and S42-IL107.

**Supplementary Figure S6.** Effects of High Ambient Temperature and *PPD-H1* on the Inflorescence Meristem Size in GP-fast and GP.

**Supplementary Figure S7.** Effects of High Ambient Temperature and *PPD-H1* on Spike Fertility and Grain Yield.

**Supplementary Figure S8.** Principle Component Analyses in the Leaf and Shoot Apex.

**Supplementary Figure S9.** Differentially Expressed Gene Identification in the Leaf and Shoot Apex.

**Supplementary Figure S10.** Function and Expression Pattern of Temperature-Responsive Genes in the Leaf and Shoot Apex.

**Supplementary Figure S11.** Expression of Known Flowering Time and Floral Development Regulators in the Leaf and Shoot Apex.

**Supplementary Figure S12.**Weighted Gene Co-Expression Network Analysis (WGCNA) of Temperature-Responsive Genes in the Shoot Apex.

**Supplementary Figure S13.** Expression of Representative Genes in the Development-Related Clusters in the Shoot Apex.

**Supplementary Figure S14.** Effects of *PPD-H1* and High Ambient Temperature on Auxin, Gibberellin, and Cytokinin Levels in the Leaf.

**Supplementary Table S1.** Effects of High Ambient Temperature on Grain Set and Rate of Spikelet Meristem and Floral Meristem Induction.

**Supplementary Table S2.** RT-qPCR Primer Used in This Study.

**Supplementary Table S3.** Unlabelled Standards and Isotopically Labelled Internal Standards (I.S.) for Phytohormone Analysis.

**Supplementary Dataset 1.** CDS Sequences of All Genotypes.

**Supplementary Dataset 2.** Transcript per Million Values of All Genes Expressed in the Leaf.

**Supplementary Dataset 3.** Transcript per Million Values of All genes Expressed in the Shoot Apex.

**Supplementary Dataset 4.** Gene Ontology Enrichment of Temperature-and *PPD-H1-*Regulated Genes in Leaf and Shoot Apex.

**Supplementary Dataset 5.** Gene List in Heatmaps.

**Supplementary Dataset 6.** Cluster-Trait Correlation.

